# The Role of *Gdf5* Regulatory Regions on Development of Hip Morphology and Susceptibility to Osteoarthritis and Dislocation

**DOI:** 10.1101/389684

**Authors:** Ata M. Kiapour, Jiaxue Cao, Mariel Young, Terence D. Capellini

**Affiliations:** Department of Orthopaedic Surgery, Boston Children’s Hospital, Harvard Medical School, Boston, Massachusetts, USA; Department of Human Evolutionary Biology, Harvard University, Cambridge, Massachusetts, USA; Farm Animal Genetic Resources Exploration and Innovation Key Laboratory of Sichuan Province, Sichuan Agricultural University, Chengdu, China; Broad Institute of Harvard and MIT, Cambridge, Massachusetts, USA

**Author notes:** **Corresponding Author:** Terence D. Capellini, PhD, Richard B. Wolf Associate Professor, Department of Human Evolutionary Biology, Harvard University, 11 Divinity Avenue, Cambridge, MA 02138.

**Keywords:** *GDF5*, Hip, Morphology, Osteoarthritis, Hip Dysplasia

## Abstract

Given *GDF5* involvement in hip development, and osteoarthritis (OA) and developmental hip dysplasia (DDH) risk, here we sought to assess the role(s) of *GDF5* and its regulatory sequence on the development of hip morphology linked to injury risk. The *brachypodism (bp)* mouse, which harbors a *Gdf5* inactivating mutation, was used to survey how *Gdf5* loss of function impacts the development of hip morphology. Two transgenic *Gdf5* reporter BAC lines were used to assess the spatiotemporal expression of *Gdf5* regulatory sequences. Each BAC line was also used to assess the functional roles of upstream and downstream sequence on hip morphology. *bp/bp* mice had shorter femora with smaller femoral heads and necks as well as larger alpha angles, smaller anterior offsets, and smaller acetabula, compared to bp/+ mice (p<0.04). Regulatory sequences downstream of *Gdf5* drove strong prenatal (E17) expression and low postnatal (6 months) expression across regions of femoral head and acetabulum. Conversely, upstream regulatory sequences drove very low expression at E17 and no detectable expression at 6 months. Importantly, downstream, but not upstream *Gdf5* regulatory sequences fully restored all the key morphologic features disrupted in *bp/bp* mice. Hip morphology is profoundly affected by *Gdf5* absence, and downstream regulatory sequences mediate its effects by controlling *Gdf5* expression during development. This downstream region contains numerous enhancers harboring risk variants related to hip OA, DDH, and dislocation. We posit that subtle alterations to morphology driven by changes in downstream regulatory sequence underlie this locus’ role in hip injury risk.

## Introduction

Several hip conditions, including osteoarthritis (OA) and developmental dysplasia of the hip (DDH), are among the most common musculoskeletal injuries. The unique morphological features of the human proximal femur and pelvic acetabulum, such as the shape and size of the femoral head, and acetabular diameter and depth, are among major contributors to hip biomechanics and stability. Changes to these morphological features often result in altered joint loading and motion, which can lead to decreased joint stability and/or increased risk of cartilage degeneration over time. These findings are supported by reports correlating the deformities of the proximal femur (i.e. cam deformity) and acetabulum (i.e. acetabular dysplasia) to increased risk of hip dislocation and OA.(1–3)

Hip morphology is initially determined during embryogenesis, when a series of flattened mesenchymal cells of the joint interzone form and delineate the boundary between the proximal femur and acetabulum. The interzone additionally provides progenitor cells that give rise to hip connective tissues (e.g. articular cartilage, ligaments, and labrum), while adjacent cells give rise to the bony elements. Over the course of late prenatal and early postnatal development, hip morphology matures and once it is mechanically loaded undergoes remodeling. Changes to this process can lead to hard and soft-tissue abnormalities, which can result in non-physiologic joint loading and ultimately increased risk of instability, injury, or degeneration.

The *Growth and Differentiation Factor 5* (*GDF5*) gene encodes a bone morphogenic protein of the TGF-ß superfamily, found only in vertebrates.(4) It is one of the key components of the biological pathways involved in pre- and postnatal development of synovial joints (e.g. hip).(5) In humans and mice, coding mutations in *GDF5* can lead to a broad spectrum of skeletal abnormalities including short stature, misregistered and malformed joints, and missing digits.(4, 6–11) More prevalent, however, is the link established in several Genome-Wide Association Studies (GWAS) between common single nucleotide polymorphisms (SNPs) spanning a 130 kb interval containing *GDF5* and OA.(12–14) This interval reflects an underlying haplotype found in hundreds of millions of people worldwide, the frequency of which has been attributable to a SNP residing within a growth plate enhancer (*GROW1*) that affects long bone size.(7) Interestingly, no variants affecting the *GDF5* or downstream *UQCC* protein coding sequences can account for the specific population-level OA associations, leading researchers to focus on linked *GDF5* 5’UTR variants. While these variants reduce transcriptional activity of the core *GDF5* promoter *in vitro,*(12) and along with other variants in strong linkage disequilibrium, correlate with reduced *GDF5* transcript levels *in vivo*,(13) they do not reside in sequence capable of specifically impacting hip morphology.(15)

Despite strong and reproducible evidence suggesting *GDF5* associations with hip OA, DDH, and dislocation.(12-14, 16-18) its mechanism of action is unknown. A recent investigation into the *cis*-regulatory architecture of the human and mouse *GDF5* locus, using a Bacterial Artificial Chromosome (BAC) scan and fine-mapping approach, revealed a number of distinct *GDF5* synovial joint regulatory enhancers.(15) Many of these enhancers reside downstream of *GDF5*, control hip expression, and harbor risk variants related to hip OA, DDH, and dislocation.(12-14, 16-18) Considering the importance of proximal femur and acetabulum morphology to hip biomechanics and stability, it is possible that the role of *GDF5* in hip OA and dislocation is mediated by alterations in hip morphology. Thus, here we sought to assess the role(s) of *GDF5* and its regulatory sequence on the development of key hip morphological features that are linked to hip injury risk

## Materials and methods

All mouse procedures were done in accordance with protocols approved by the Harvard University Institutional Animal Care and Use Committee (IACUC; protocol # 13-04-161).

### Mouse Strains

To study how *Gdf5* loss-of-function impacts hip morphology, the BALB/cJ *bp*^3J^ strain was used, in which a CG dinucleotide is replaced by a single T at position 876 within the *Gdf5* coding sequence(4, 15) truncating the entire mature sequence and rendering the allele non-functional. Mice with two copies of this allele (*bp/bp*) lack *Gdf5* and exhibit the classic brachypodism phenotype, while mice with one copy (*bp/+*) have no overt developmental abnormalities.(6) Since the typical breeding strategy for this line generates *bp/bp* and *bp/+* animals, we used *bp/+* mice as controls in this study as they are indistinguishable from wild type mice. To study the impact of broad regulatory regions on *Gdf5* function, two transgenic FVB-NJ strains were also used (Upstream BAC and Downstream BAC),(6, 15) each expressing *Gdf5* and *lacZ* from a BAC transgene containing mouse *Gdf5* and ~110-kb of upstream or downstream flanking sequence. Accordingly, each BAC contained an IRES-β-Geo cassette within the *Gdf5* 3’UTR, allowing for bicistronic transcription of *Gdf5* and *lacZ*. For rescue experiments, stable *Upstream BAC* and *Downstream BAC* lines were separately crossed to *bp/bp* mice. *Upstream BAC;bp/+* and *Downstream BAC;bp/+* were next back-crossed to *bp/bp* mice, and progeny were genotyped for the *lacZ* transgene and the *bp* allele in separate PCR reactions as described.(15)

### Skeletal Preparations and Staining

Euthanized adult 8-week old mice were subjected to cartilage and bone skeletal staining protocols using Alcian blue and Alizarin red, respectively. In brief, euthanized mice were skinned, eviscerated, and muscle surrounding most long bones and axial tissues was removed. The specimens were then ethanol dehydrated, acetone treated, Alcian blue/Alizarin red S stained, and finally cleared using 1% KOH/glycerol.(6)

### Micro-Computed Tomography (MicroCT) Imaging and Morphology Assessment

To quantify phenotypic defects related to *Gdf5* loss-of-function as well as to gauge the influence that regulatory sequences have on hip morphology, right femora and pelves of twenty 8-week old mice (N=5 per genotype) were scanned using high-resolution MicroCT (µCT40, SCANCO Medical AG, Brüttisellen, Switzerland). DICOM images were exported for measurements of several clinically relevant morphological features of the femur and pelvis in Osirix MD v7.5 (Pixemo SARL, Bernex, Switzerland) using established protocols (Figure S1).(19–22) The measurer (AMK) was blinded to specimens’ genotype. Data normality was assessed by Shapiro-Wilk’s test in SPSS (IBM Corp., Armink, NY). Normally distributed data were compared using Analysis of Variance (ANOVA) with a post hoc Tukey correction for multiple comparisons. Non-normally distributed data were compared between the groups using Kruskal-Wallis test with a Benjamini Hochberg post hoc correction for multiple comparisons (Prism, GraphPad Software Inc., La Jolla, CA). P-values are two-sided and the statistical significance was assessed at alpha = 0.05.

## Results

### Assessment of the Role of *Gdf5* in the Development of Hip Morphology

Examination of microCT imaged skeletal preparations of *Gdf5* null (*bp/bp*) and control mice reveals that qualitatively, 8-week old *bp/bp* mice possess a normal looking hip joint, with all of the prominent morphological features of the proximal femur and acetabulum (Figure 1). However, detailed quantitative analyses reveal significant alterations to several hip features, especially those linked to risk of injury. Compared to *bp/+* mice, *bp/bp* mice had shorter femora (p=0.014), smaller femoral head diameters (p=0.001), smaller femoral head offsets (p<0.001), shorter femoral neck lengths (p=0.009), and smaller femoral neck diameters (p<0.001). Moreover, *bp/bp* mice had larger alpha angles (p=0.038), smaller anterior offsets (p=0.037), smaller acetabular diameters (p<0.001) and shallower acetabular depths (p=0.004) (Figure 2). Conversely, there are no significant differences in valgus cut angles (p=0.087), neck shaft angles (p=0.220), femoral head tilt angles (p=0.465), anteversion angles (p=0.506) and pelvis lengths (p=0.789) between genotypes. The mean (SD) of all quantified indices along with the P-values for all pairwise comparisons are presented in Table S1.

**Figure 1:**
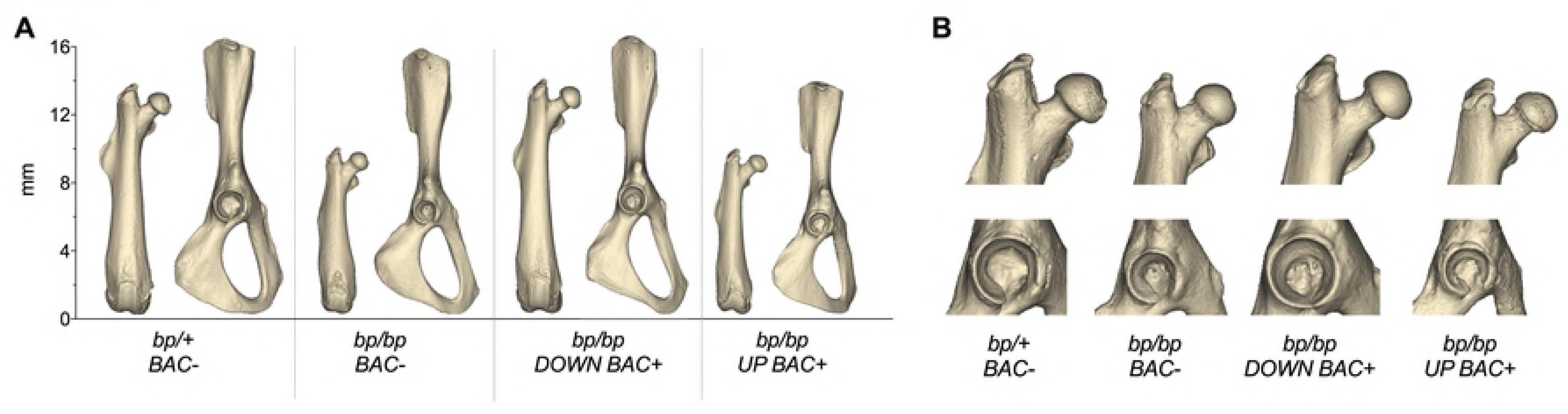
Reconstructed three-dimensional models are shown for (A) femur and pelvis and (B) close up views of proximal femur and acetabulum from a representative specimen of each genotype. BAC, Bacterial Artificial Chromosome; *bp*, brachypodism; *DOWN BAC+*, *Downstream BAC*; *UP BAC+*, *Upstream BAC*.

**Figure 2:**
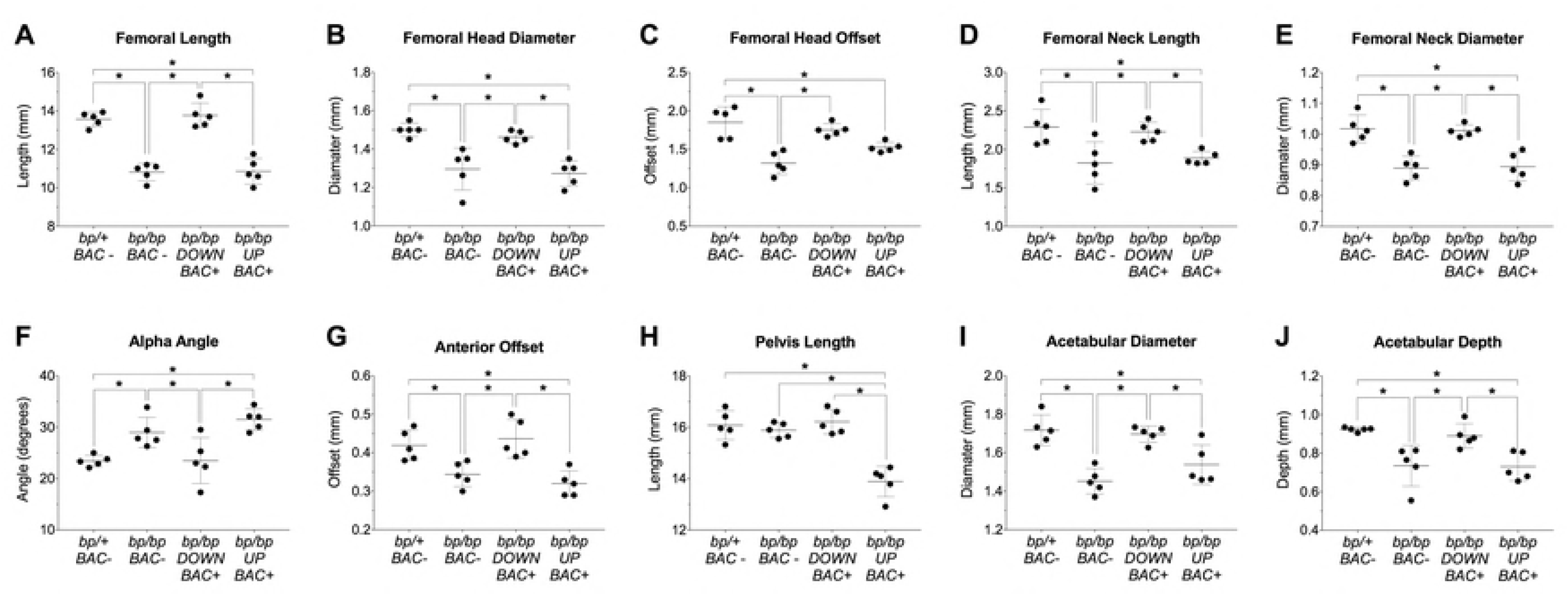
Quantitative morphological analysis of the femur and pelvis in the *bp/bp* and *bp/bp+BAC* rescue experiments. Group differences in (A) femoral length (* p<0.02), (B) femoral head diameter (* p<0.01), (C) femoral head offset (* p<0.01), (D) femoral neck length (* p<0.03), (E) femoral neck diameter (* p<0.01), (F) alpha angle (* p<0.04), (G) anterior offset (* p<0.04), (H) pelvis length (* p<0.05), (I) acetabular diameter (* p<0.01), and (J) acetabular depth (* p<0.02). Mean ± SD (n=5/group). BAC, Bacterial Artificial Chromosome; *bp*, brachypodism; *DOWN BAC+*, *Downstream BAC*; *UP BAC+*, *Upstream BAC*.

### Late Prenatal and Postnatal Expression of *Gdf5* Regulatory Domains

Given the conservation of the mouse and human *GDF5* gene as well as its regulatory sequence function,(7, 15) we next assessed in late fetal and postnatal mouse hips the expression patterns of two large *Upstream BAC* and *Downstream BAC* clones, each expressing *Gdf5* and *lacZ* from a BAC transgene containing mouse *Gdf5* and ~110-kb of upstream or downstream flanking sequence. During prenatal stages (at E17), the *Downstream BAC* drove expression ubiquitously throughout the hip joint with strong expression throughout the proximal femur and acetabulum (Figure 3); specifically, across the proximal femoral head articular cartilage, femoral neck, trochanters and intertrochanteric zones, and proximal chondrocyte growth zone. Within the acetabulum, the *Downstream BAC* drove expression across the labrum, the acetabular surface and rim, the lunate surface, ligaments, as well as the surrounding chondrocytes in the growth zone formed between the ilium, ischium, and pubis. In contrast, at E17, the *Upstream BAC* drove expression at a much lower level and only across the femoral head articular cartilage as well as partially in the labrum and acetabulum proper (Figure 3; black arrowheads). By 6 months postnatal, a much-reduced expression pattern was observed for the *Downstream BAC* construct: expression was observed only at the most proximal, anterior region of the femoral head articular cartilage as well as along the superior lip of the labrum and acetabulum rim (Figure 3; white arrowheads), locations of major joint loadings in mice. In contrast, at 6 months postnatal, no detectable *lacZ* expression was observed in the proximal femur and acetabulum using the *Upstream BAC* construct (Figure 3). Detailed expression patterns of each *BAC* are presented in Supplementary Table S2.

**Figure 3:**
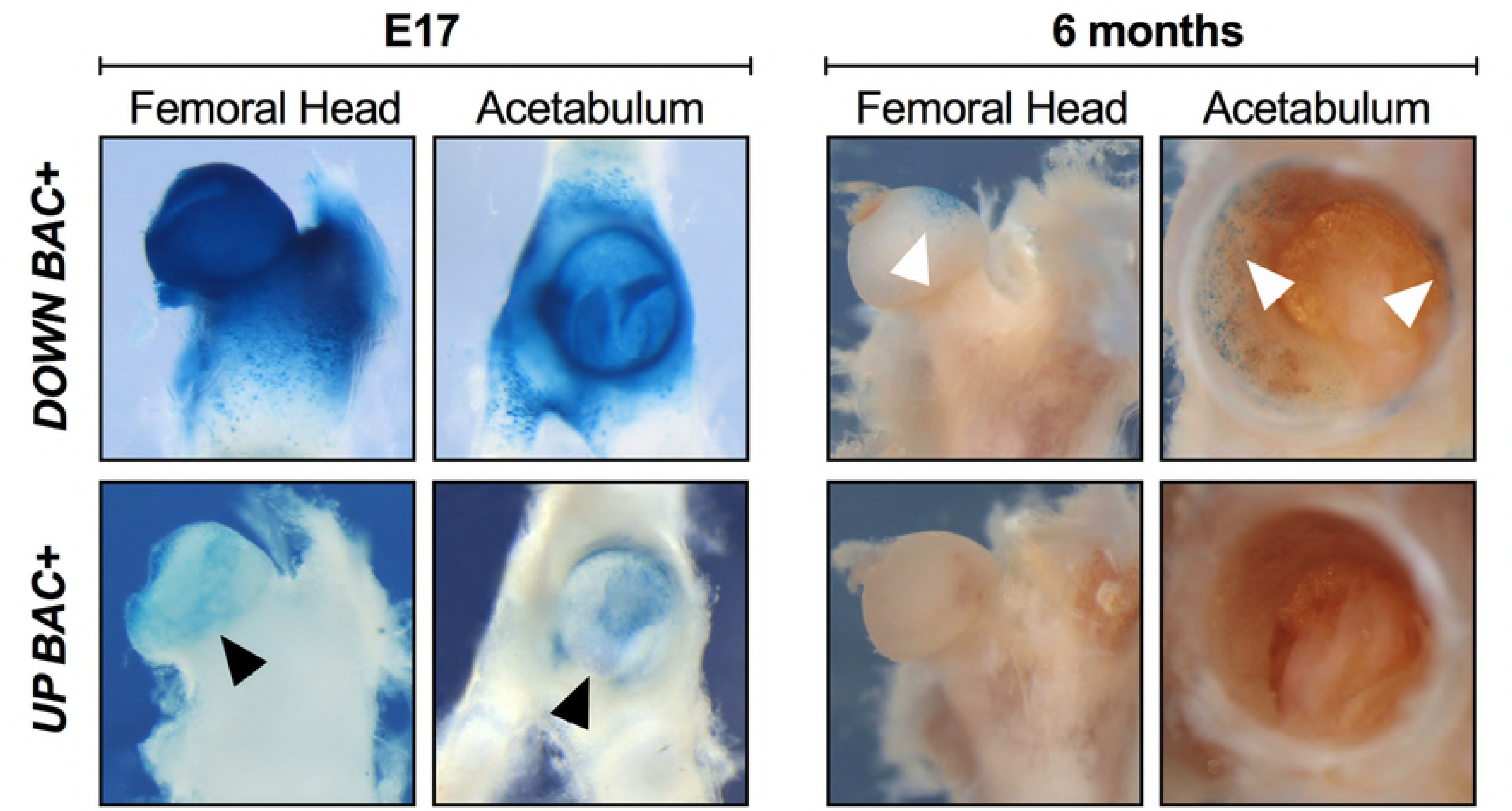
Comparison of *Upstream* and *Downstream BAC* expression in late prenatal hip development and in the postnatal hip. X-gal-stained hips from *Downstream BAC* (*DOWN BAC+*) and *Upstream BAC* (*UP BAC+*) mice at E17 and 6 months are shown. At E17, the *Downstream BAC* drove a strong, ubiquitous expression throughout the hip joint covering the proximal femur and acetabulum, whereas the *Upstream BAC* drove expression at a much lower level and only across the femoral head articular cartilage as well as partially in the labrum and acetabulum proper (black arrowheads). At 6 months, the D*ownstream BAC* construct drove a much-reduced expression pattern, with expression present only at the most proximal, anterior region of the femoral head articular cartilage as well as along the superior lip of the labrum and acetabulum rim (white arrowheads). No detectable expression was observed using the *Upstream BAC* construct at 6 months. Scale bars: E17 = 0.5 mm; 6 month (Proximal Femur) = 2 mm; 6 month (Acetabulum) = 0.75 mm.

### Assessment of *Gdf5* Regulatory Sequence in Shaping Hip Morphology

Given the dynamic expression patterns by both the *Upstream* and *Downstream BAC* constructs, and prior findings indicating their influence on long bone and knee morphology,(6, 7, 15) we more deeply explored the regulatory control that each regulatory sequence block has on hip morphology. Taking advantage of our bicistronic *Gdf5-lacZ* construct design, we introduced a copy of each BAC transgene onto the *bp/bp* mouse background and gauged phenotypic rescue at 8 weeks using microCT and morphometric techniques. Transgenic expression of *Gdf5* by the *Downstream BAC* restored the femoral length as well as all quantified morphological features of the proximal femur and acetabulum that were disrupted in the *bp/bp* mouse (Figure 2 and Table S1; p>0.8 for all *bp/+* vs. *bp/bp* Down BAC+ comparisons). In contrast, *Gdf5* expression through the *Upstream BAC* failed to restore femur length and the dysmorphic morphology of the proximal femur and acetabulum of *bp/bp* mice (Figure 2 and Table S1; p<0.03 for all *bp/+* vs. *bp/bp* Up BAC+ comparisons).

## Discussion

In this study, we investigated the influence that *Gdf5* and its broad regulatory domains have on hip morphology, in the context of shape parameters relevant to OA, DDH, and dislocation. We revealed that: (1) *Gdf5* loss results in substantial dysmorphologies to the proximal femur and acetabulum in line with known features involved in hip instability, developmental dislocation, injury, and adult onset OA; (2) Compared to upstream regulatory sequences, downstream regulatory sequences of *Gdf5/GDF5* drive stronger, more ubiquitous prenatal hip expression and uniquely control postnatal hip expression; (3) This regulatory structure is functionally important as downstream sequences were capable of restoring *Gdf5* loss-of-function phenotypes to normal, a situation not observed when using the upstream region. These latter two findings have important ramifications to how we understand *GDF5* and its association with OA and hip dysplasia/dislocation.

Indeed, this is the first study to not only demonstrate a direct link between *Gdf5* loss-of-function and hip dysmorphology in mice but to demonstrate that loss of *Gdf5* influences proximal femur and acetabular morphology in a manner concordant with known hip joint features that are clinically relevant to hip ailments. Interestingly, unlike prior observations that the *bp/bp* knee joint is completely dysmorphic,(6) gross morphology of its hip joint appeared relatively normal. However, our use of quantitative imaging approach revealed that fine-grained features of both the proximal femur and acetabulum were significantly affected. The anatomical alterations that we observed have been either directly linked to hip OA risk in humans (e.g. femoral head diameter, femoral neck length, and femoral neck diameter) or previously reported in patients with femoroacetabular impingement (FAI), DDH, and recurrent hip dislocation, which are proven risk factors for hip OA.(1–3) Abnormal development of these features can compromise hip stability (e.g. hip dislocation) and/or result in non-physiologic joint loading, leading to excessive cartilage wear and degeneration (e.g. OA) overtime.

Importantly, there is ample evidence revealing the importance of non-coding regulatory variants in the *GDF5* locus as potentially underlying hip dysplasia’s and degenerative diseases.(12-14, 16-18) These types of regulatory mutations often have less pleiotropic effects on phenotypes. For example, recent reports have shown substantial variations in these hip morphologic features between subjects with progressive hip OA and matched controls,(2, 23, 24) and candidate association and GWAS studies have repeatedly revealed the association of the *GDF5* locus with OA, including that of the hip. In these studies, a large 130 kb haplotype underlies risk of OA at the locus.(12–14) Likewise, several studies further demonstrate that variants far downstream of *GDF5*, within the same risk haplotype are also associated with DDH.(16–18) In both cases, the underlying causal variants likely reside within developmental enhancers in relevant regulatory domains.

Accordingly, here we have revealed that a broad downstream regulatory domain, which completely spans the *GDF5* hip OA and DDH risk interval, possesses the functional ability to significantly influence hip morphology. We specifically identified that the *Downstream BAC* allele restored all abnormal *bp/bp* measures of proximal femoral and acetabular morphology to normal control levels, whereas the regulatory sequences upstream of *Gdf5*/*GDF5* had little impact on phenotypic rescue. Within this downstream regulatory domain, at least four published (*GROW1, R3, R4*, and *R5*) and three unpublished *GDF5* regulatory sequences exist (data not shown), several of which (*GROW1*, *R4*, and two others) drive hip expression, and harbor risk variants.(7, 15)

While these findings do shed light on the regulatory control of Gdf5 during hip formation, this study has some scientific limitations. First, given the nature of breeding the *bp/bp* line, all genotypic comparisons are made to heterozygous (*bp/+*) controls, which have been extensively reported upon as having no overt limb abnormalities or evidence of OA or hip defects up through six months of age. Likewise, we only concentrated on male mice given the opportunistic numbers acquired through breeding, though, at the *GDF5* locus sex-specific effects in OA and DDH risk have not been reported.

In summary, this work helps to elucidate the role(s) that *Gdf5* and its regulatory domains have in hip development and provides an important developmental context for subsequent studies that address the impact of human *GDF5* genetic variants on a range of hip injuries including OA. In light of our findings and given that genetic variants in the *GDF5* association interval, as well as hip morphology, are risk factors for OA and DDH, we propose a developmental-genetic model in which risk-associated variants in downstream regulatory sequences influence local *GDF5* expression levels in developing hip structures, leading to subtle alterations in hip morphology that, in turn, predispose the joint to injury and subsequent degeneration. It is also possible that such variants act to decrease *GDF5* levels in matured adult hips and that this may also influence OA risk by impairing homeostasis in healthy joints or by accelerating degeneration due to injury. The development of a conditional allele for disrupting *Gdf5* expression at specific pre- and postnatal time-points and spatial domains, along with the targeted deletion of its hip joint enhancers will be essential for teasing apart potential roles of *Gdf5* in hip development and/or homeostasis. Finally, an extensive functional interrogation of the downstream regulatory and its variants should prove fruitful for identifying causative variants underlying hip OA and DDH at this locus.

## Acknowledgements

The authors would like to thank David Kingsley at Stanford University, Hao Chen at Genentech, Michele Schoor at Miltenyi Biotec, Hari Reddi at the UC Davis Center for Tissue Regeneration and Repair, and Daniel Brooks and Mary Bouxsein from the Imaging and Biomechanical Testing Core of the Center for Skeletal Research (NIH P30 AR066261) at the Massachusetts General Hospital. This work was funded in part by the Milton Fund and Dean’s Competitive Fund of Harvard University, NIH (NIAMS) grants to TDC (R01AR070139) and AMK (R01AR065462), and the Department of Orthopaedic Surgery at Boston Children’s Hospital.

## Funding

This work was funded in part by the Milton Fund (TDC) and Dean’s Competitive Fund of Harvard University (TDC), NIH (NIAMS) grants to TDC (R01AR070139) and AMK (R01AR065462), and the Department of Orthopaedic Surgery at Boston Children’s Hospital (AMK). The views expressed are those of the authors and not necessarily those of the funding agencies. None of the Sponsors had any involvement in the design, collection, analysis or interpretation of the data and no role in writing or submitting this manuscript for publication.

## Competing Interest

Authors have nothing to disclose.

## Data Availability

All relevant data are within the paper and its supporting Information files.

## Supplementary materials

**Figure S1:** Quantified anatomical indices of the femur and pelvis. Measurements were conducted on microCT images acquired at 12 µm3 isotropic voxel size, 70 kVp peak x-ray tube intensity, 114 mA x-ray tube current, and 200 ms integration time. Femoral length (FL), femoral head diameter (FHD), femoral head offset (FHO), femoral neck length (FNL), femoral neck diameter (FND), valgus cut angle (VCA), neck shaft angle (NSA), femoral head tilt angle (FHTA), alpha angle (AA), anterior offset (AO), pelvis length (PL), acetabular diameter (ADI) and acetabular depth (ADE).

**Table S1:** Differences in key morphologic features of the hip joint at 8 weeks. All outcome measures were defined as continuous variables (N=5 per genotype). Normally distributed data were compared using Analysis of Variance (ANOVA) with a post hoc Tukey correction for multiple comparisons. Non-normally distributed data were compared between the groups using Kruskal-Wallis test with a Benjamini Hochberg post hoc correction for multiple comparisons. P values are two-sided and the statistical significance was assessed at alpha = 0.05 for all the comparisons.

**Table S2:** BAC prenatal and postnatal expression patterns.

**AUTHOR CONTRIBUTIONS**
T.D.C. conceived and oversaw the project. T.D.C helped perform mouse rescue experiments. J.C. and M.Y. performed mouse breeding and collected bp and BAC-LacZ tissue samples. A.M.K. performed all morphometric and statistical analyses on bp and BAC-rescue mice. A.M.K and T.D.C. wrote the manuscript with input from all other authors.

